# Skeletal muscle DNMT3A plays a necessary role in endurance exercise by regulating oxidative capacity of red muscles

**DOI:** 10.1101/2020.05.18.102400

**Authors:** Sneha Damal Villivalam, Dongjoo You, Scott M. Ebert, Jinse Kim, Han Xiao, Hector H. Palacios, Christopher M. Adams, Sona Kang

## Abstract

Exercise interventions alter the DNA methylation profile in skeletal muscle, yet little is known about the role of the DNA methylation machinery in exercise capacity. In this study, we found that in oxidative red muscle, DNMT3A expression increases greatly following a bout of endurance exercise. Mice lacking *Dnmt3a* in skeletal muscle fibers had reduced tolerance to endurance exercise, accompanied by reduced oxidative capacity and reduced mitochondrial counts. Moreover, during exercise, the knockout muscles overproduced reactive oxygen species (ROS), which are major contributors to muscle dysfunction. In mechanistic terms, we demonstrated that *Aldh1l1* is a key target of repression by DNMT3A in red muscles. DNMT3A directly regulated the Aldh1l1 transcription by binding to the *Aldh1l1* promoter region and altering DNA methylation and histone modification. Enforcing ALDH1L1 expression, leading to elevated NADPH, led to overproduction of ROS by the NADPH oxidase complex (NOX) in myotubes, ultimately resulting in mitochondrial defects. Moreover, both genetic inhibition of ALDH1L1 and pharmacological inhibition of NOX rescued oxidative stress and mitochondrial decline in *Dnmt3a*-deficient myotubes, confirming the essential role of ALDH1L1-dependent ROS generation as a downstream effector of DNMT3A loss of function. Together, our results reveal that DNMT3A in skeletal muscle plays a pivotal role in endurance exercise by controlling intracellular oxidative stress.

## Introduction

Physical exercise maintains health, prevents disease, and even acts as medicine for a wide range of non-communicable diseases by improving whole-body metabolism and inducing beneficial adaptations in multiple tissues throughout the body [1,2]. Exercise can be classified broadly into two types, endurance and resistance. Endurance exercise (*e.g*., marathon running and swimming), also called aerobic exercise, is generally characterized by high frequency, long duration, and low power output. By contrast, resistance exercise (*e.g*., body building and throwing events), which is based on strength, is characterized by activities of low frequency, high resistance, high intensity, and short duration [3,4].

Endurance exercise primarily relies on slow-twitch muscles, such as the soleus, which are also called red muscles due to their high myoglobin content [5]. Red muscles contain more mitochondria than fast-twitch (white) muscles and are more aerobic; consequently, they rely on oxidative metabolism, which favors fatty acids as a fuel source [5]. Accordingly, endurance exercise stimulates mitochondrial biogenesis and the expression of genes involved in mitochondrial respiration and β-oxidation of free fatty acids, driving a phenotypic adaptation toward a more oxidative phenotype [6]. Reduced oxidative potential due to mitochondrial decline or dysfunction is often associated with exercise intolerance and muscle disorders [7–10].

Reactive oxygen species (ROS), including oxygen-derived molecules such as hydrogen peroxide (H_2_O_2_) and free radicals such as superoxide (•O_2 −_) and hydroxyl radical (HO•), cause oxidative stress [11,12]. A moderate increase in skeletal muscle ROS production in the acute phase of exercise is thought to activate signaling pathways that lead to cellular adaptation, thereby protecting against future stress [13–17]. However, excessive ROS can oxidatively damage macromolecules including DNA, lipids, and proteins, as well as modify cellular redox status and cellular functions; consequently, ROS elevation is also associated with pathophysiological states of muscle and contractile dysfunction [13–16]. Mitochondria make a large contribution to ROS production at rest, but not during muscle contraction [13,17–19]. The majority of ROS produced during contraction arise from non-mitochondrial sources, such as NADPH oxidase (NOX), located in the microtubules [16,20,21]. The redox-mediated crosstalk between NOX and mitochondria exacerbates ROS production and disrupts redox homeostasis [21–24]. For example, NOX-derived ROS promote the opening of mitochondrial ATP-sensitive K^+^ channels [23–25]. The resultant potassium influx into the matrix lowers the mitochondrial membrane potential, which causes mitochondrial swelling, opening of permeability transition pores, and elevated ROS production [23–25]. In addition, NOX-derived ROS causes leakage of Ca^2+^ from the sarcoplasmic reticulum or entry of extracellular Ca^2+^, resulting in mitochondrial Ca^2+^ overload and mitochondrial ROS emission, which ultimately results in muscle fatigue and dysfunction [13,17–19,26–28].

Exercise significantly alters the DNA methylation profile of skeletal muscle [29–37]. DNA methylation, a reversible epigenetic mark that usually occurs on a cytosine residue followed by a guanine (CpG), is mediated by a member of the DNA methyltransferase (DNMT) family [38]. Methylation prevents the binding of transcriptional machinery that requires interaction with cytosine, usually resulting in transcriptional silencing [39]. Acute and chronic forms of exercise induce both hyper- and hypo-CpG methylation of target loci [29–37], and some of these modifications are inversely correlated with gene expression [29,34,37,40]. For example, a single bout of aerobic endurance exercise in human subjects transiently induces promoter hypomethylation in important mitochondria-related transcripts (e.g., *PPARGC1A, PDK4, TFAM*, and *PPARD*), followed by an increase in their expression [29]. Despite the obvious link between altered DNA methylation and exercise, few studies have investigated the roles of individual DNMTs in exercise and muscle adaptations.

Here, we report that skeletal muscle DNMT3A is a critical epigenetic modulator of endurance exercise. In this study, we discovered that DNMT3A in soleus, a red muscle, increases after a bout of endurance exercise. In mice, muscle-specific *Dnmt3a* deletion led to exercise intolerance and reduced muscle strength under various exercise regimens and was accompanied by increased signs of myopathy. Knockout (KO) soleus muscle exhibited a dramatic reduction in oxidative capacity and decline in mitochondrial count, causing the tissue to favor non-oxidative glycolysis rather than fatty acid oxidation. We identified *Aldh1l1* as a key direct target of repression by DNMT3A in soleus muscle. Overexpression of ALDH1L1 was sufficient to recapitulate *Dnmt3a* KO–mediated mitochondrial dysfunction, and also caused oxidative stress by promoting accumulation of NADPH and thereby increasing NOX activity. Furthermore, *Aldh1l1* KO or pharmacological inhibition of NOX rescued mitochondrial decline and oxidative stress caused by *Dnmt3a* deficiency. Together, our results provide novel insights into the epigenetic regulation of the muscle response to exercise and reveal a surprising molecular target that is important for sustaining endurance exercise.

## Results

### DNMT3A level in the red soleus muscle increases after endurance exercise

DNMT1 is the major enzyme involved in maintenance of the DNA methylation pattern following DNA replication, whereas DNMT3A and DNMT3B are primarily responsible for de novo DNA methylation [38]. Hence, we postulated that de novo DNMTs might be more important for adaptive responses to environmental changes. To begin to characterize the role of de novo DNMTs in endurance exercise, we examined their expression patterns in soleus, extensor digitorum longus (EDL), and gastrocnemius (GA) muscles, which are red, white, and mixed muscles, at rest and after a bout of endurance exercise, in C57BL/6J wild-type mice. We also measured the mRNA expression of PPARγ-coactivator 1α (*Ppargc1a*), which is induced by exercise in skeletal muscle [29,41,42]. *Ppargc1a* mRNA levels increased after 50 minutes of treadmill running, especially in the soleus and GA (**Supplemental Fig. 1A**). Strikingly, *Dnmt3a* mRNA levels also increased by ∼3 fold after exercise in both soleus and GA but not in EDL (**Supplemental Fig. 1B-D**). By contrast, exercise caused no significant changes in *Dnmt3b* transcript levels in any muscle types (**Supplemental Figs. 1B-D**). The increase in the level of DNMT3A after a bout of exercise was confirmed by western blotting (**Supplemental Fig. 1A**).

### Muscle-specific *Dnmt3a* deficiency decreases the capacity for endurance exercise

The increase in DNMT3A expression in soleus (red) and GA (mixed) but not in EDL (white) suggested that DNMT3A plays an important role in endurance exercise, which primarily relies on red muscles that are largely dependent on aerobic respiration. To address this question, we used muscle creatine kinase (MCK)-Cre to generate muscle-specific knockout mice (MCK-*Dnmt3a* KO) (**Supplemental Figs. 2A, B)**. MCK-*Dnmt3a* KO mice were viable, exhibited normal growth and fertility, and displayed no significant differences in body weight, body composition, energy balance, or muscle mass relative to wild-type littermates harboring two floxed alleles without Cre (WT) (**Supplemental Fig. 3)**.

To assess tolerance to endurance exercise, we employed two different regimens: (1) a low-intensity regimen (**Supplemental Fig. 4A**) that tested the ability to run steadily at relatively low speed (12 m/min) for an initial 40 min, followed by a gradual increase in speed until exhaustion [43], and (2) a high-intensity regimen (**Supplemental Fig. 4B**) that rapidly increased the running speed (6 m/min and increased by 2 m/min every 5 min) to a maximal pace of 30 m/min, which persisted until exhaustion [43,44]. The first regimen preferentially tests fatty acid oxidation capacity, whereas the latter focuses on glucose usage efficiency [43,44]. The running capacity of the MCK-*Dnmt3a* KO mice was greatly impaired: both distance and duration were reduced by 30–40% under both the low- and high-intensity regimens (**Figs. 1A, B**).

**Fig. 1.**
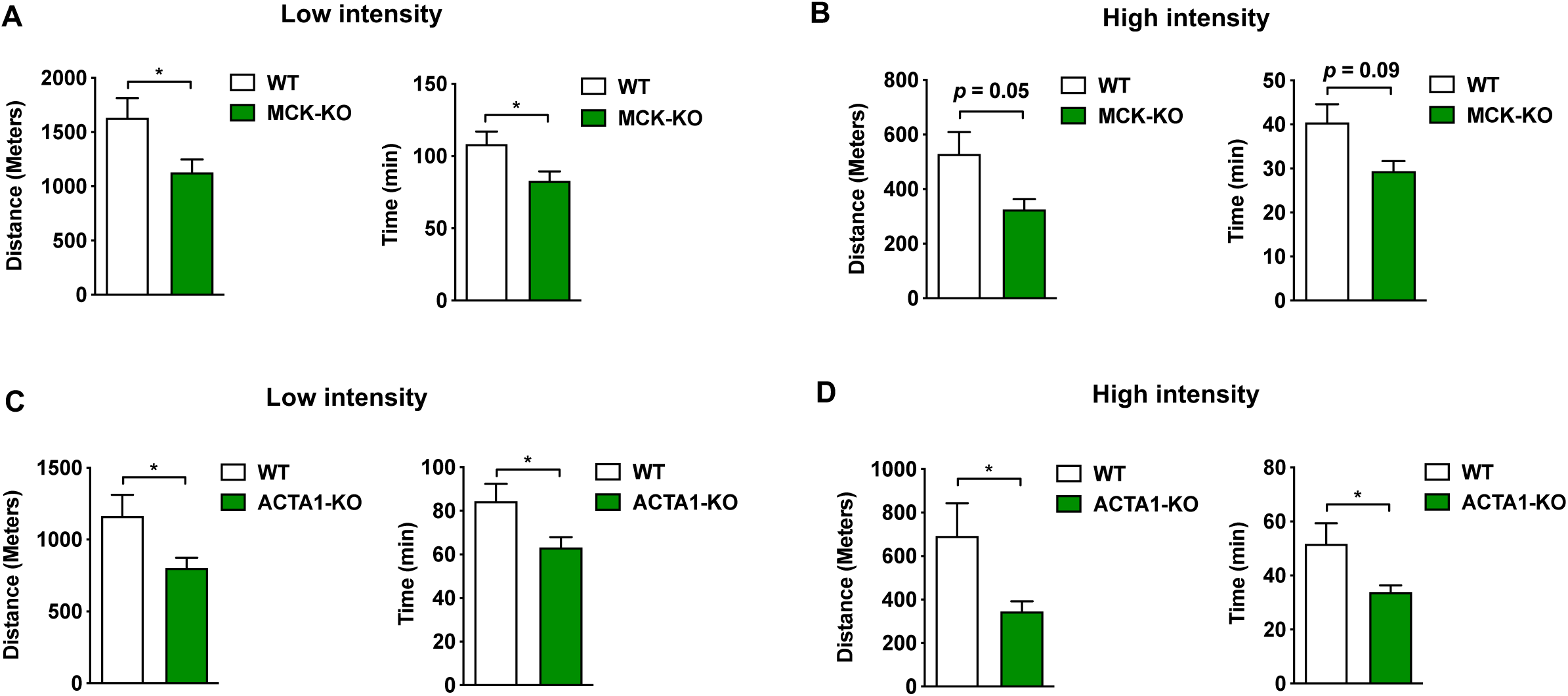
Muscle-specific *Dnmt3a* KO mice display a reduced tolerance to endurance exercise. (**A, B**) The low intensity (**A**) and high intensity (**B**) regiment was conducted in MCK-*Dnmt3a* KO and WT mice (n = 5 mice, p<0.05, Student’s *t*-test, mean ±s.e.m.). (**C, D**) The low intensity (**C**) and high intensity (**D**) regiment was conducted in ACTA1-*Dnmt3a* KO and WT mice (n = 7, p<0.05, Student’s *t*-test, mean ±s.e.m.).

Cardiac muscle [45,46] plays an important role in exercise by maintaining blood flow to muscles [47]. Because the Cre recombinase in the MCK-Cre line is also expressed in cardiac muscle [45,46], we had to consider a cardiac contribution to the exercise incapacity of the KO mice. To investigate this possibility, we generated an independent line of muscle-specific *Dnmt3a* KO mice, ACTA1-*Dnmt3a* KO, using human alpha-skeletal actin (ACTA1-Cre-ERT), which drives Cre expression in skeletal muscle in a tamoxifen-dependent manner, without expression in cardiac muscle (**Supplemental Figs. 5A, B**). To achieve muscle-specific *Dnmt3a* KO in this model, we injected tamoxifen (50 mg/kg per day × 5 days, intraperitoneal [i.p.]) into 8-week-old *Dnmt3a*_*f*/*f*_/ACTA1-Cre^+^ mice; their tamoxifen-treated Cre^-^ littermates were used as controls. After 2 weeks of tamoxifen washing, we subjected the mice to the treadmill regimens to test their exercise capacity. As with MCK-*Dnmt3a* KO mice, ACTA1-*Dnmt3a* KO mice displayed an exercise intolerance of a similar magnitude in both regimens (**Figs. 1C, D**), ruling out a cardiac contribution to their exercise incapacity.

During endurance exercise, metabolism shifts its fuel source from carbohydrates to fatty acids in order to spare glucose for the brain and sustain muscle energy [7,48]. Hence, we investigated whether reduced exercise intolerance was associated with altered fuel utilization. In both KO models, blood glucose levels at rest did not differ from those of controls (**Supplemental Figs. 6A, D**), indicating that the reduced exercise capacity was not simply due to hypoglycemia. Following exercise, however, KO mice exhibited subtly reduced plasma glucose levels (**Supplemental Fig. 6A, D**) and elevated lactate levels (**Supplemental Fig. 6B, E**) relative to WT mice, implying that KO mice favor non-oxidative glycolysis. On the other hand, free fatty acid levels were reduced following exercise relative to controls (**Supplemental Figs. 6C, F**). Notably, voluntary physical activity of the MCK-*Dnmt3a* KO mice, as assessed using a CLAMS unit (**Supplemental Figs. 3E, F**), was not altered, eliminating the possibility that reduced exercise capacity was due to impaired voluntary movement.

### *Dnmt3a*-KO muscle has reduced oxidative capacity and diminished mitochondrial content

High oxidative potential and mitochondrial content of skeletal muscle is critical for supporting endurance exercise. Succinate dehydrogenase (SDH), located in the inner membrane of the mitochondrion, is responsible for oxidizing succinate to fumarate in the citric acid cycle [49]. Hence, we performed SDH staining to distinguish between oxidative and less-oxidative muscles. In KO animals, both soleus and GA muscles exhibited a dramatic (50%) decline in oxidative potential (**Figs. 3A–D**). Consistent with this observation, mitochondrial copy number was reduced by 43% and 47% in the red soleus and mixed GA muscles, respectively (**Fig. 3E**), but was unchanged in the white EDL muscle (**Fig. 3E**). The magnitude of the decrease in mitochondrial content was similar in the soleus and GA muscles, accounting for their reduced oxidative capacity. Consistent with decreased mitochondrial DNA amount, reduced oxygen consumption rate was noted in KO soleus muscles (**Figs. 3F, G**). Further, we noted that Dnmt3a knock-down in mature L6 myotubes has the same effects on mitochondrial counts and oxygen consumption rates (**Figs. 3I-J**).

**Fig. 2.**
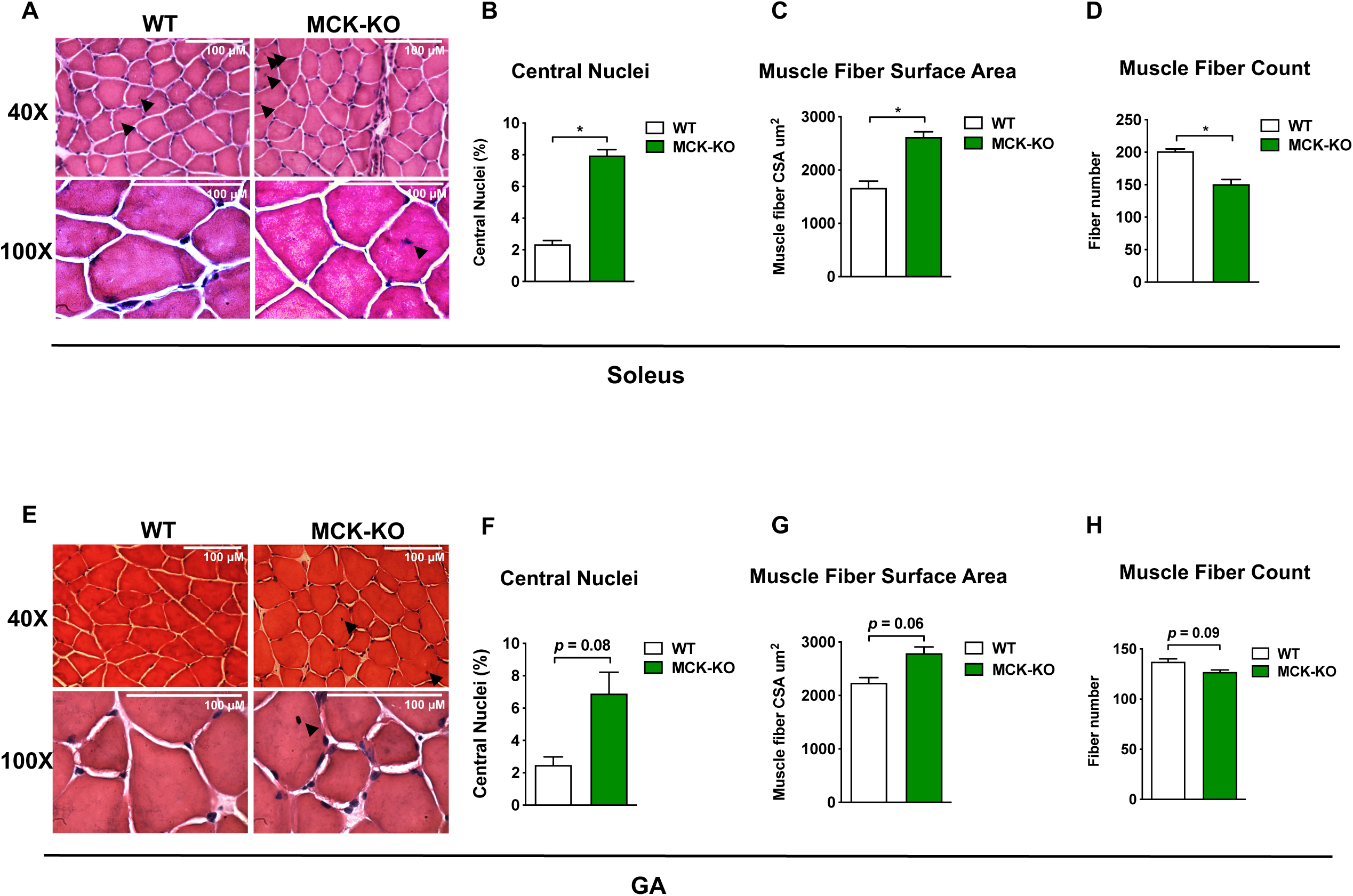
Muscle-specific *Dnmt3a* KO mice display increased muscle damage and myopathy following exercise. (**A**) The H&E staining of KO and WT soleus muscle after a bout of low intensity treadmill running (top 40X, bottom 100X magnifications). Black arrow indicates centralized nuclei. (**B**) The percentage of myofibers with centralized nuclei was determined by manual counting 100–150 myofibers in 20X magnification. Muscle fiber surface area (**C**) and muscle fiber count (**D**) was measured in WT and KO soleus muscle by ImageJ using 100–150 myofibers in 20X magnification. (Cross sectional area (CSA) (n = 2 mice, p<0.05, Student’s *t*-test, mean ±s.e.m.). (**E-H**) The analogous set of data is shown with WT and KO GA muscles.

**Fig. 3.**
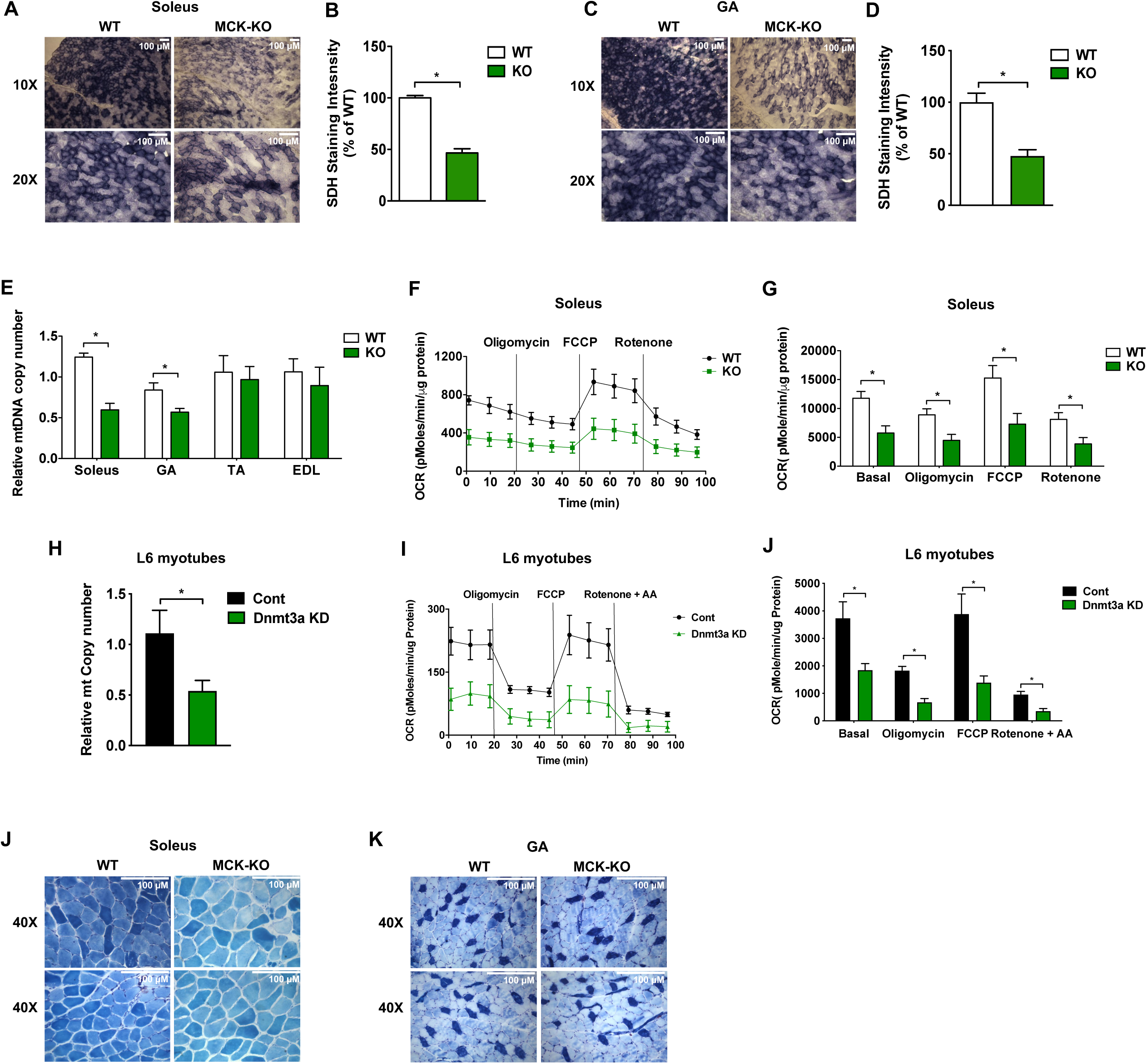
*Dnmt3a*-KO soleus muscle displays a decreased oxidative capacity with a reduced mitochondrial content. (**A-D**) Succinate dehydrogenase staining was performed in WT and KO soleus (**A, B**) and GA (**C, D**) muscles after a bout of low-intensity exercise for 50 min (10X, 20X magnifications), and the staining intensity was quantified using ImageJ (n = 2, p < 0.05, Student’s t-test, mean ± s.e.m.). (**E**) Genomic DNA was extracted from various muscle types from WT and KO mice after a bout of low-intensity exercise for 50 min. Mitochondrial DNA copy-number was calculated from the ratio of mitochondria-encoded COXII to nuclear-encoded cyclophilin A (n = 6, p < 0.05, Student’s t-test, mean ± s.e.m.). (**F, G**) Mitochondrial respiration was measured in WT and KO soleus tissue after a bout of low-intensity exercise for 50 min under basal conditions and in response to 1.5 mM oligomycin (complex V inhibitor), 4 mM FCCP (uncoupler), or 2 mM rotenone (complex I inhibitor) (*n* = 3, p < 0.05, Student’s *t*-test, mean ± s.e.m.). (**H-J**) L6 myotubes were transduced with lentiviral expression plasmids for Flag-ALDH1L1 and GFP and mitochondrial DNA copy number and mitochondrial respiration was was measured from these cells (n = 6, p<0.05, Student’s *t*-test, mean ±s.e.m.). (**K, L**) ATPase staining was conducted in soleus (**K**) and GA muscles (**L**), (type I: dark blue; type IIa: lightest blue; type IIb: light blue, 40X magnification).

Myosin heavy chains (MHC) are responsible for the contractility of muscle [50,51] and are thus responsible for how aerobic a muscle is. Red muscles like soleus are predominantly enriched in MHCI fibers and some IIa fibers, whose biophysical attributes confer a slow-twitch phenotype, making the muscle more aerobic. By contrast, white muscles like EDL are enriched in the faster MHCIIb fibers that are responsible for the fast twitch property. To determine whether the change in oxidative potential is due to a change in fiber type distribution in KO muscle, we performed chromatographic ATPase staining, which is more permissive for myosin heavy chain (MHC) I ATPase than for MHCII ATPase activity [52]. Overall intensity of ATPase staining was diminished in KO soleus muscle, but unchanged in KO GA muscle (**Figs. 3K, L**), suggesting an overall reduction in the oxidative capacity of both muscle fiber types in soleus, rather than fiber-type skewing. Indeed, we observed no discernable change in the expression of muscle fiber type–specific MHC isoforms between WT and KO muscles (**Supplemental Fig. 7**). Collectively, our data suggest that DNMT3A is required for the full oxidative capacity of skeletal muscle and is not associated with fiber type determination.

### Gene profiling studies identify muscle-specific DNMT3A target genes

We and others have shown that DNMT3A regulates biological processes by regulating non-overlapping sets of cell type–specific target genes [53–55]. To elucidate the underlying molecular basis by which *Dnmt3a* regulates exercise capacity, we performed RNA-Seq on WT and KO soleus muscle. The transcriptome profiles revealed 26 genes that were upregulated, and 4 that were downregulated in *Dnmt3a*-deficient soleus muscle (**Figs. 4A, B**). These data are consistent with the expectation that DNMT3A works primarily as a repressor of gene expression.

**Fig. 4.**
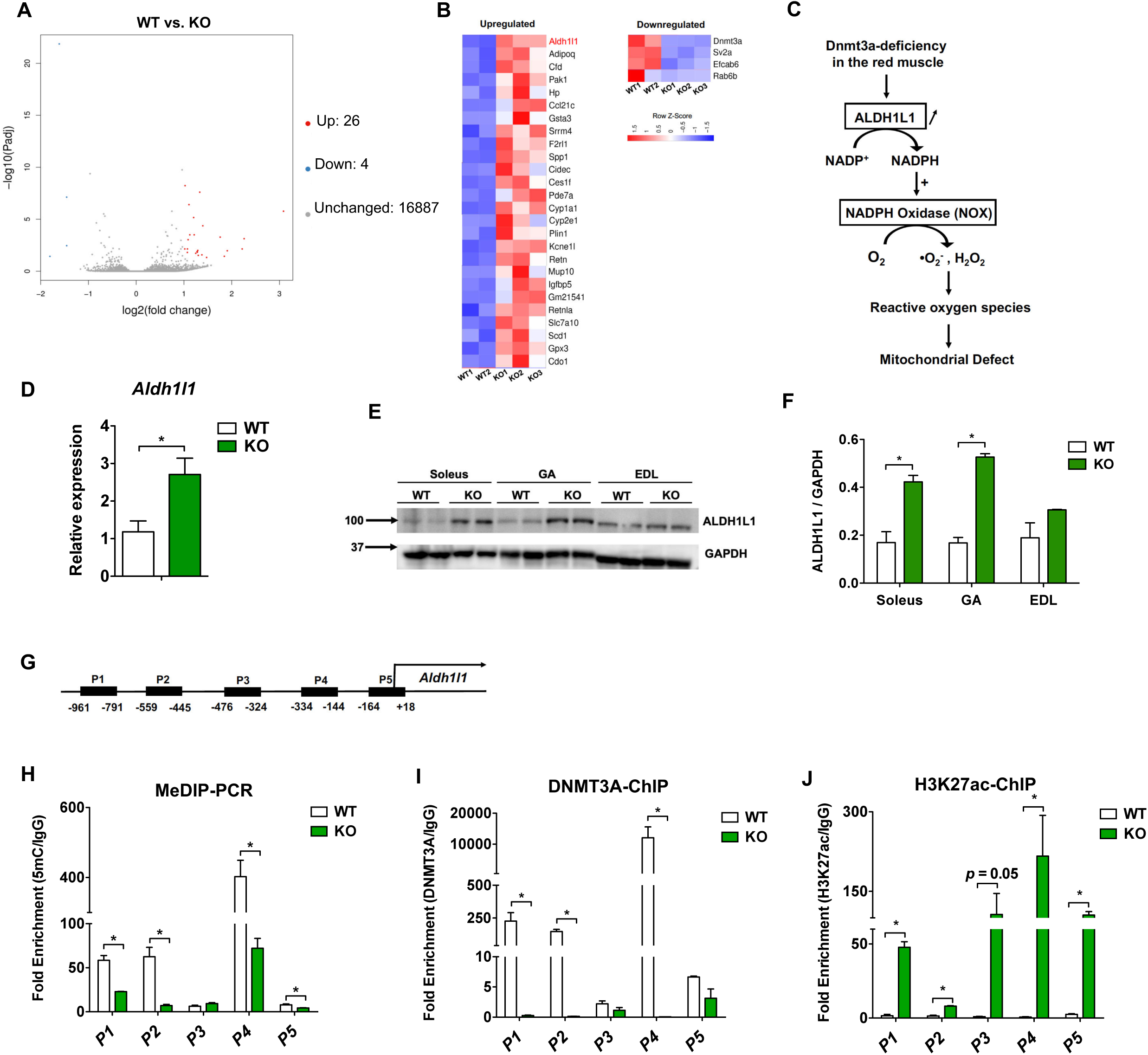
Transcriptome analysis identifies *Aldh1l1* as a key target gene of DNMT3A in the soleus muscle. (**A**) RNA-Seq was performed in WT and KO soleus muscle at rest. Volcano plot shows differentially regulated genes (Up; Log_2_FoldChange≧1, P<0.05, Down; Log_2_FoldChange≧-1, P<0.05, Unchanged Log_2_FoldChange <1, P≧ 0.05). (**B**) The heat map shows differentially expressed genes in KO soleus muscle (FDR<0.05, p < 0.05). (**C**) Proposed model for ROS regulation during loss of *Dnmt3a*. Loss of DNMT3A increases ALDH1L1 expression, thus leading to the increase in NADPH levels and increased activity of NADPH oxidase, leading to increased ROS levels. The increased oxidative stress contributes to mitochondrial dysfunction and muscle fatigue. (**D**) Q-PCR validation of *Aldh1l1* (*n* = 6, p < 0.05, Student’s *t*-test, mean ± s.e.m.). (**E, F**) ALDH1L1 protein expression and the quantification in WT vs. KO soleus muscles. (**G**) The map of CpG-rich promoter regions of *Aldh1l1* and MeDIP and ChIP primers (P1-P5) that cover the CpG rich regions. The numbers correspond to the position from the transcriptional start site of *Aldh1l1*. (**H**) MeDIP-qPCR was performed in WT and KO soleus muscle to assess differential methylation using primer sets from **F** (*n* = 3, p < 0.05, Student’s *t*-test, mean ± s.e.m.). (**I**) DNMT3A ChIP-PCR was conducted in WT and KO soleus muscles (*n* = 3, p < 0.05, Student’s *t*-test, mean ± s.e.m.). (**J**) H3K27ac ChIP-PCR was conducted in WT and KO soleus muscles (*n* = 3, p < 0.05, Student’s *t*-test, mean ± s.e.m.).

Because a relatively small number of genes were differentially regulated in the KO tissue, we did not identify any specific biological pathways enriched among the DNMT3A target genes. However, genes involved in the detoxification of reactive oxygen species (e.g., Sod1 and Gpx3) were upregulated in KO muscle, implying that the level of oxidative stress is elevated in KO muscles. Indeed, that was the case: In both MCK- and ACTA1-*Dnmt3a* KO muscles, ROS levels were significantly elevated at rest relative to controls, and the difference grew after exercise (**Supplemental Figs. 8A, B**). Importantly, *Dnmt3a* knockdown in L6 myotubes was sufficient to recapitulate the increase in ROS levels in Dnmt3a deficiency (**Supplemental Fig. 8C**), suggesting a cell-autonomous role for Dnmt3a in this context.

Next, we sought to identify the targets responsible for the increase in ROS production. As a candidate, we turned our attention to *Aldh1l1*, which encodes aldehyde dehydrogenase 1 family member L1 (ALDH1L1), a cytosolic enzyme involved in folate and one-carbon metabolism. Specifically, ALDH1L1 oxidizes 10-formyltetrahydrofolate to tetrahydrofolate, producing NADPH as a byproduct [56] (**Fig. 4C**). Notably in this regard, NADPH plays a dual role in the regulation of oxidative stress. On the one hand, it is a reducing agent for glutathione, thioredoxins, peroxiredoxins, and glutathione peroxidases, which neutralize ROS [57]. On the other hand, it contributes to ROS generation through the activity of the NADPH oxidase complex (NOX) (**Fig. 4C**), which is located within the sarcoplasmic reticulum, transverse tubules, and sarcolemma in skeletal muscle fibers [22]. We hypothesized that ALDH1L1-mediated NADPH production feeds into NOX, thereby increasing intracellular ROS in *Dnmt3a* KO muscle.

Prior to functional study, we studied the regulation of *Aldh1l1* by DNMT3A. First, we measured the levels of *Aldh1l1* mRNA and ALDH1L1 protein were elevated in KO muscle tissues (**Figs. 4D–F**). Importantly, ALDH1L1 expression was elevated only in the red soleus and mixed GA muscles, but not in the white EDL muscle (**Figs. 4E, F**), suggesting that DNMT3A-mediated repression of *Aldh1l1* is red muscle–specific. To determine whether *Aldh1l1* is a direct target of DNMT3A, we conducted methylated DNA immunoprecipitation (MeDIP)-qPCR analysis of the CpG rich *Aldh1l1* promoter regions (**Fig. 4G)**. In KO soleus muscle, DNA methylation was greatly reduced in four of the five regions we tested, including the CpG island (P4), (**Fig. 4H**). *In vivo* ChIP assay confirmed strong enrichment of DNMT3A at those differentially methylated regions (**Fig. 4I**). In gene regulation, the DNA methylation and histone regulation machineries often engage in crosstalk [58]. Therefore, we conducted a ChIP assay for H3K27ac, a histone modification marker of active promoters and enhancers, and detected strong signals at *Aldh1l1* promoter regions in KO soleus tissues (**Fig. 4J**). Collectively, these data demonstrate that DNMT3A directly regulates gene expression of *Aldh1l1* by modifying the epigenetic profile at its regulatory regions.

### ALDH1L1 drives the increase in ROS level and mitochondrial dysfunction in KO soleus muscle

To determine whether elevated ALDH1L1 expression is responsible for NADPH-dependent generation of ROS, we compared NADPH levels in WT and KO muscle. Indeed, NADPH levels were significantly increased in KO muscle (**Fig. 5A**). To assess whether this resulted in increased flux into NOX, we measured the NADPH consumption rate [59,60] and detected elevated NOX activity in KO tissues (**Fig. 5B**). Next, to evaluate the functional significance of ALDH1L1, we performed an ALDH1L1 gain-of-function study in L6 rat myotubes in the presence and absence of apocynin, a specific NOX inhibitor [61]. Overexpression of ALDH1L1 had no effect on myogenesis (**Supplemental Fig. 9**), but remarkably, it was sufficient to recapitulate the redox changes in *Dnmt3a*-KO muscles, including the increases in the levels of NADPH (**Fig. 5C**) and ROS (**Figs. 5D, E**). More strikingly, ALDH1L1-overexpressing myotubes had reduced mitochondrial copy number (**Fig. 5F)** and oxygen consumption relative to controls (**Figs. 5G, H**).

**Fig. 5.**
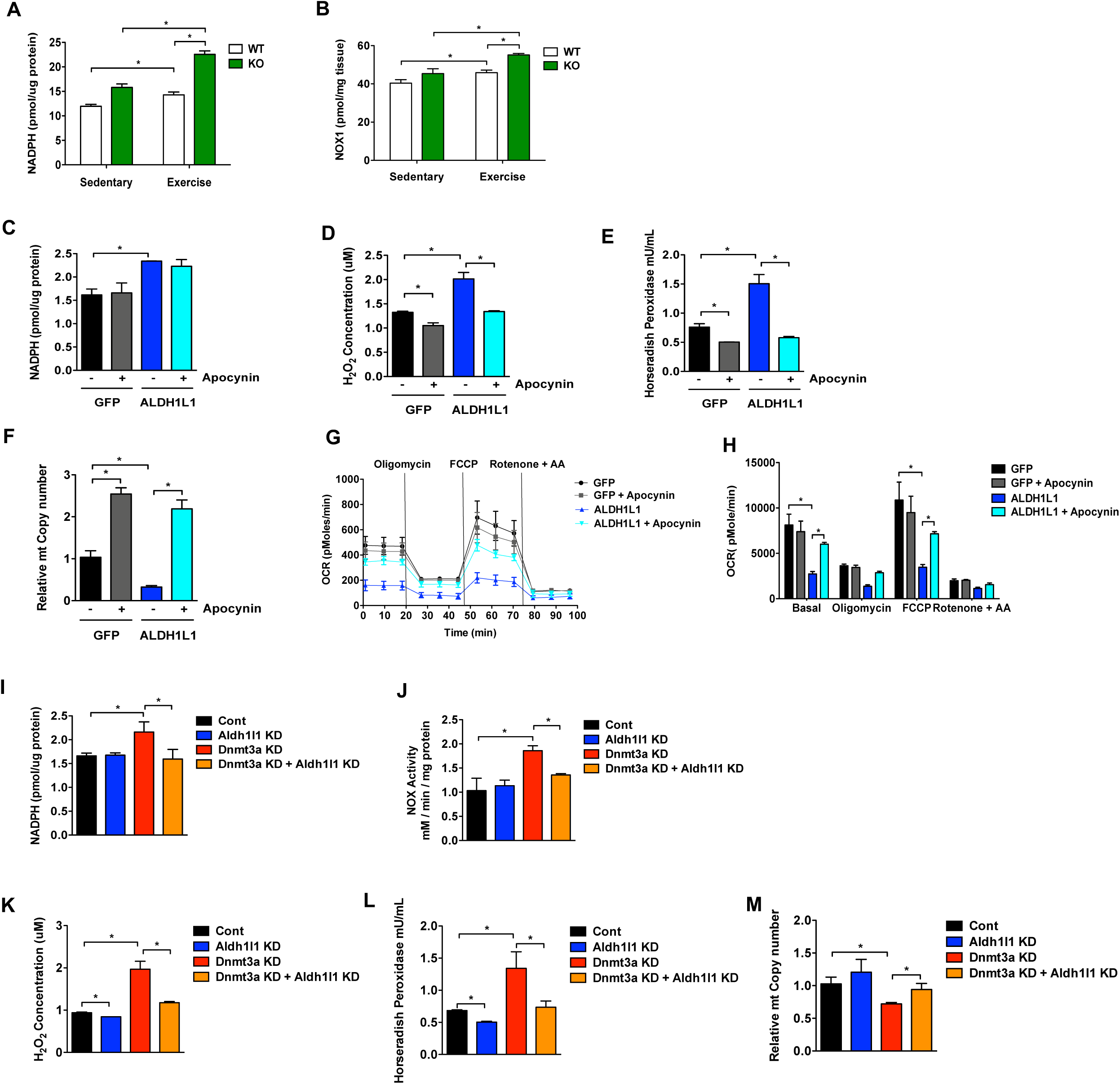
ALDH1L1 contributes to the oxidative stress and mitochondrial defect in loss of *Dnmt3a*. NADPH levels (**A**) and NOX (**B**) activity was measured in WT and KO muscles at rest and after a bout of low-intensity exercise for 50 min (*n* = 6, p < 0.05, Student’s *t*-test, mean ± s.e.m.). (**C-H**) L6 myotubes were transduced with lentiviral expression plasmids for Flag-ALDH1L1 and GFP. NADPH levels (**C**), ROS levels (**D, E**), Mitochondrial DNA copy number (**F**), Mitochondrial respiration (**G, H**) were measured from these cells in the presence and absence of NADPH oxidase inhibitor apocynin (*n* = 3, p < 0.05, 2-way ANOVA, mean ± s.e.m.). (**I-M**) Single and double knockdowns of *Dnmt3a* and *Aldh1l1* in L6 myotubes were achieved by lentiviral transduction. NADPH levels (**I**) and NOX activity (**J**) were measured in single and double knockdowns of *Dnmt3a* and *Aldh1l1* in L6 myotubes. (*n* = 6, p < 0.05, Student’s *t*-test, mean ± s.e.m.). (**K, L**) ROS levels were measured in single and double knockdowns of *Dnmt3a* and *Aldh1l1* in L6 myotubes. (*n* = 6, p < 0.05, Student’s *t*-test, mean ± s.e.m.). (**M**) Mitochondrial DNA copy number was measured in single and double knockdowns of *Dnmt3a* and *Aldh1l1* in L6 myotubes. (*n* = 6, p < 0.05, 2-way ANOVA, mean ± s.e.m.).

Importantly, all of these changes associated with ALDH1L1 overexpression were largely reversed by treatment with apocynin (**Figs. 5C-H**). To prove that ALDH1L1 is essential for the phenotype of the *Dnmt3a* loss-of-function model, we knocked out *Aldh1l1* in *Dnmt3a* knockdown L6 myotubes, which resulted in dramatic rescue of the oxidative stress and mitochondrial decline caused by *Dnmt3a* deficiency (**Figs. 5I–M**).

## Discussion

Physical inactivity contributes to lifestyle-related diseases, including obesity, type 2 diabetes, cardiovascular diseases, and age-related muscle wasting [1,2,62,63]. Although the benefits of exercise have long been known, its underlying regulatory mechanisms are still not fully understood. Exercise significantly alters the DNA methylation profile of skeletal muscle [29–37]. For example, a single bout of aerobic exercise in human subjects transiently induces promoter DNA hypomethylation in important mitochondria-related transcripts (e.g., *PPARGC1A, PDK4, TFAM*, and *PPARD*), followed by an increase in their expression [29]. In addition, in healthy adult human subjects, moderate-intensity exercise results in hypermethylation of *FABP3* and *COX4L1*, which is negatively correlated with their expression [36,64]. Collectively, these data suggest that changes in promoter methylation are associated with alterations in gene expression following aerobic exercise. By the same token, genome-wide studies have reported that both acute and chronic exercise interventions produce profound changes in CpG methylation [29–37]. Some of the alterations are concordant with changes in gene expression [29,34,37,40]. Despite the apparent connection between altered DNA methylation and exercise, the underlying roles of DNMTs in exercise performance remain unclear. In this study, using two independent genetic models of muscle-specific *Dnmt3a*-KO mice, we showed that DNMT3A plays an essential role in endurance exercise.

Skeletal muscle mitochondria are highly dynamic organelles that exhibit remarkable plasticity, adapting their content, structure, and metabolism in response to a variety of physiological and pathophysiological stresses including exercise, disuse, and aging [65–67]. Exercise training increases mitochondrial biogenesis to satisfy elevated energy requirements by increasing oxidative capacity to ensure optimal ATP supply; this has the consequence of favoring lipid metabolism [68–70]. Thus, exercise represents a viable therapy, with the potential to reverse the impairment of mitochondrial function associated with diseases such as muscular dystrophy, atrophy, type 2 diabetes, and aging-related sarcopenia [1,2,71–81].

A key link between exercise and control of mitochondrial biogenesis was revealed by the observation that PGC-1α expression is transiently induced in skeletal muscle following an acute bout of exercise [82]. Since that discovery, a great deal of research effort has been devoted to elucidating the role of PGC1A in skeletal muscle mitochondria biology and exercise. For example, transgenic expression of PGC1A increases mitochondrial content and function, increases the abundance of oxidative type I muscle fibers, and decreases muscle fatigue. However, loss of PGC1A has only a mild effect on exercise capacity and does not alter fiber-type composition in muscle [83] or affect training-induced increases in the expression of genes involved in oxidative phosphorylation [84]. This suggests that PGC1A is sufficient, but not necessary, to mediate metabolic adaptations in response to exercise. In this study, we found that DNMT3A is required for mitochondria and metabolic adaptations in red skeletal muscle. Because we did not detect obvious changes in the expression of PGC1A (not shown), we speculate that the role of DNMT3A in the regulation of mitochondrial function and mass is likely to be PGC1A-independent.

Overproduction of ROS induced by unaccustomed, exhaustive exercise training or other stresses can lead to oxidative stress–related tissue damage and reduced contractility [85,86]. NADPH oxidases are major contributors to ROS production [87,88]. Previous work showed that physical stretching can increase the activity of NADPH oxidase, especially NOX2, to produce ROS in microtubule-dependent process [89]. That study described mechanotransduction-dependent activation of NADPH in cardiac muscle; here, by contrast, we identified ALDH1L1-dependent activation of NADPH oxidase in skeletal muscles. Our finding that inhibition of NADPH oxidase rescued both oxidative stress and decline in mitochondrial content raises the possibility of repurposing inhibitors to improve exercise trainability.

Although we focused on *Aldh1l1*, DNMT3A has other target genes worthy of additional attention. Indeed, candidate genes revealed by our transcriptomics analysis suggest that several secretory factors involved in fibrosis and inflammation are up-regulated in Dnmt3a-deficient muscles. For example, *Spp1* encodes osteopontin (OPN), also referred to as bone sialoprotein 1 (BSP-1), which contributes to fibrosis, inflammation, and collagen deposition in aged animals and models of muscle dystrophy, e.g., *Mdx* mice [90,91]. Additionally, we noted that *Ccl21c* and *Ccl21d*, both of which encode the proinflammatory chemokine CCL21, were upregulated in Dnmt3a-deficient muscles. Notably in this regard, both OPN and CCL21 promote migration of immune cells such as macrophages, neutrophils, or T cells [91–93]. Therefore, it is conceivable that elevated expression of these factors may slow down recovery from myopathy in Dnmt3a-deficient muscle following exercise or muscle damage. Future studies should address these possibilities.

Here, we highlighted the surprising role of DNMT3A in endurance exercise, skeletal muscle mitochondrial biology, and energy utilization. In mechanistic terms, we revealed that ALDH1L1 serves as a novel molecular link that contributes to oxidative stress and mitochondrial dysfunction following the loss of *Dnmt3a* in red muscle. This is of great importance from the standpoint of exercise physiology, as physical activity is strongly encouraged as a key strategy for preventing and treating a wide range of human diseases. Understanding the epigenetic and molecular basis of exercise tolerance will help us to address several critical health issues that arise due to lack of exercise.

## Materials and Methods

### Animals

Animal Care Mice were maintained under a 12-hr light /12-hr dark cycle at constant temperature (23°C) with free access to food and water. All mice were extensively back-crossed onto a C57Bl/6J background. All animal work was approved by UC Berkeley and the University of IOWA ACUC. *In vivo* assays were done with 7- to 20-week-old littermate male mice.

### Measurement of exercise capacity

All mice were acclimated to the treadmill prior to the exercise test session. For each session, food was removed 2 hr before exercise. Acclimation began at a low speed of 5 to 8 meters per minute (m/min) for a total of 10 min on Day 1 and was increased to 5 to 10 m/min for a total of 10 min on Day 2. The experiments were performed on Day 3. For the low intensity treadmill test, the treadmill began at a rate of 12 m/min for 40 min. After 40 min, the treadmill speed was increased at a rate of 1 m/min every 10 min for a total of 30 min, and then increased at the rate of 1 m/min every 5 min until the mice were exhausted. The high intensity treadmill test was conducted on the same open-field six-lane treadmill set at a 10% incline. Following a 5-min 0 m/min acclimation period, the speed was raised to 6 m/min and increased by 2 m/min every 5 min to a maximal pace of 30 m/min until exhaustion. Mice were considered exhausted when they were unable to respond to continued prodding with a soft brush.

### Grip strength assessment

Grip strength was measured either with a wire or inverted wire mesh grid. For wire hanging, each mouse was placed with their forelimbs on a 2 mm wire that was tied from end to end. The hang time was measured for each mouse until they dropped down. For the string test, each mouse was placed with their forelimbs on a 2 mm wire that was tied from end to end. The time taken for mice to hold on to the wire with their hindlimbs was noted. Mice that failed to hold on with hind limbs were not included in the calculation. For the cage top test, mice were placed on top of an elevated wire mesh grid, the screen was inverted, and the animals were timed until they let go of the grid.

### RNA extraction and quantitative PCR

Total RNA was extracted from tissues using TRIzol reagent according to the manufacturer’s instructions. cDNA was reverse transcribed from 1 μg of RNA using the cDNA Reverse Transcription Kit (Applied Biosystems). Quantitative PCR (qPCR) was performed with SYBR green qPCR master mix (AccuPower 2X, Bioneer) using a CFX96 Touch (Bio Rad). The relative amount of mRNA normalized to cyclophilin B was calculated using the delta–delta method. Primer sequences are listed in **Supplemental Table 1**.

### Western blot analysis and antibodies

Whole-cell protein lysates were prepared according to the manufacturer’s protocol using RIPA lysis buffer and protease inhibitor cocktail. Proteins were size fractionated by SDS-PAGE and then transferred to polyvinylidene difluoride membranes. After blocking with 5% nonfat dried milk in TBS-Tween (0.25%), the membranes were incubated with the appropriate primary antibodies against antiDNMT3A antibodies (#2160S). The loading control included anti-GAPDH (#2118) and anti-Histone H3 (#14269). Immunoblots were quantified by ImageJ.

### Immunoprecipitation

For immunoprecipitation, tissues were lysed with 300 μl lysis buffer, containing 50 mM Tris-HCl (pH7.5), 150 mM NaCl, 1% Triton X-100, and protease inhibitor cocktail for 30 min on rotation. The lysates were centrifuged at 13,000 rpm for 10 min at 4°C, then supernatants were incubated with antiDNMT3A antibodies (#2160S) overnight then with protein A/G PLUS-Agarose (#sc-2003) at 4°C for 2 hr. The beads were washed four times with lysis buffer. Bound proteins were eluted by boiling in SDS sample buffer and processed for western blot.

### Genomic DNA extraction

Snap-frozen tissues were treated with proteinase K overnight at 56°C. DNA was extracted by standard phenol chloroform–ethanol precipitation and eluted in DNase-free water.

### Histological analysis of skeletal and cardiac muscle fibers

Harvested skeletal muscles were immediately embedded in T.F.M. compound (Tissue-Tek) and snap frozen using a Stand-Alone Gentle Jane (Instrumedics Inc.). We then prepared 10 μm sections from the muscle mid-belly using a Leica cryostat. Hematoxylin and eosin (H&E) staining was performed following fixation in ice-cold zinc formalin (Anatech Ltd. #175) for 60 min. Image analysis was performed using ImageJ software. For ATPase stains, slow-type fibers were dark, whereas fast-type fibers were lightly stained following preincubation at pH 4.35. Fiber type was identified on the basis of color for each myofiber. For Sirius Red staining, cardiac sections were then incubated in Bouin’s’ solution (5% acetic acid, 9% formaldehyde, and 0.9% picric acid) at room temperature for 1 hr. Next, after washing, slides were incubated in 0.1% Fast Green (Fisher, F-99) for 3 min, then in 0.1% Sirius Red (Direct red 80, Sigma, 0-03035) for 2 min. After three washes, slides were dehydrated with ethanol and xylene based on standard procedures.

### Mitochondrial DNA copy number

Mitochondrial DNA (mtDNA) copy number was used as a marker for mitochondrial density. Briefly, after isolating genomic DNA from muscle tissues, mtDNA was quantified using quantitative RT-PCR. The mitochondrial DNA copy number was calculated from the ratio of COX II (mitochondrial-encoded gene)/cyclophilin A (nuclear-encoded gene).

### Measurement of plasma parameters

Mice were either kept at resting or allowed to run for 50 min on the low-intensity regime. Following this, blood was collected from the tail tip for WT and MCK-*Dnmt3a* KO mice from both groups. Blood plasma was separated by centrifugation for 15 min at 4,000 rpm. Plasma nonesterified fatty acids were measured using the NEFA-HR2 Assay Kit (Wako Diagnostics). Lactate levels in the plasma were measured using a Lactate Colorimetric Assay Kit (Abcam).

### Reactive oxygen species generation

Intracellular accumulation of reactive oxygen species (ROS) was monitored using OxiSelect™ Hydrogen Peroxide/Peroxidase Assay Kit (Cell Biolabs, Inc., San Diego, CA). To investigate ROS generation, the muscle tissues were homogenized in 1× assay buffer provided from the kit. The lysates were assayed according to the manufacturer’s procedure.

### RNA-Seq library generation and analysis

RNA samples were extracted using the RNeasy Mini kit (Qiagen, 74104), and the quality of total RNA was assessed by the 2100 Bioanalyzer (Agilent) and agarose gel electrophoresis. Libraries were prepared using the BGI Library Preparation Kit, and sequencing was performed on the BGISEQ (BGI, China). RNA-Seq reads were aligned to the UCSC mm10 genome using HISAT2 (Hierarchical Indexing for Spliced Alignment of Transcripts [94]), and mapping was done using Bowtie2 [94]. Differentially regulated genes were calculated using DEseq2 [95].

### Measurement of NADPH

The NADPH measurement was performed on GA/Soleus extracts using the NADP/NADPH assay kit (Cat#K347, BioVision) according to the manufacturer’s instructions. Briefly, ∼20 mg samples were extracted in 400 µL of the given extraction buffer, and 50 µL was processed following instructions of the kit. OD450 measurements were made on a plate-reader (SpectraMAX i3 Plate reader) at 25°C, and the data was calculated using a standard curve.

### Measurement of NADPH oxidase activity

NOX activity was measured by accessing oxidation of NADPH through Continuous Spectrophotometric Rate Determination [59,60]. Briefly, samples of tissue or cells were extracted in 500 µL and 200µL of potassium phosphate buffer, respectively. The tissues and cells were first homogenized and then sonicated. The homogenate was centrifuged at top speed for 10 min, and the supernatant was used for reading. Oxidation of NADPH was monitored at 340 nm on the SpectraMAX i3 Plate Reader at 30°C [59,60].

### MeDIP-qPCR

Genomic DNA was sheared using a Covaris S220 to an average of 200–600 bp. 600 ng of denatured DNA was incubated with 2 mg of 5-methylcytosine (5-mC) monoclonal antibody ([33D3] Diagenode, Cat No # C15200081) in IP buffer (0.1% SDS, 1 Triton X-100, 2 mM EDTA, 20 mM Tris-HCl pH 8.1, 150 mM NaCl) for 1 h at 4°C on a rotating wheel. Antibody-bound DNA was collected with 20 ml of protein A/G PLUS-Agarose (#sc-2003) for 1 h at 4°C on a rotating wheel and successively washed three times with IP buffer (0.1% SDS, 1 Triton X-100, 2 mM EDTA, 20 mM Tris-HCl pH 8.1, 150 mM NaCl). DNA was recovered in 100 ml of digestion buffer (50 mM Tris pH 8.0, 0.5% SDS, 35 mg proteinase K) and incubated overnight at 65°C. Recovered DNA was used for qPCR analysis. Primers for MeDIP-qPCR studies are listed in **Supplementary Table 1**. All data were normalized to input.

### Cell culture

L6 rat myoblasts were obtained from UCSF and were cultured in Dulbecco’s modified Eagle’s medium (DMEM) supplemented with 10% fetal bovine serum. Differentiation was carried out in Dulbecco’s modified Eagle’s medium (DMEM) supplemented with 10% horse serum, 100 U/ml penicillin, and 100 μg/ml streptomycin. Myocytes were maintained in a humidified incubator under an atmosphere of 5% CO_2_ at 37°C. To generate lentivirus particles, lentiviral constructs were co-transfected with pMD2.G- and psPAX2-expressing plasmids into 293T cells. After 48 h, the virus-containing supernatant was collected, filtered through 0.45-mm filters, and added to mature L6 myotubes for 48 h along with 8 mg/ml polybrene. Transduction efficiency was determined by comparing to cells transduced in parallel with a GFP-expressing lentivirus, and myotubes were treated with 100 μM Apocynin (Tocris, Cat. No. 4663) for 24 hours.

### Metabolic cages

Indirect calorimetry and food intake, as well as locomotor activity, were measured using the Comprehensive Lab Animal Monitoring System (CLAMS) (Columbus Instruments). The calorimetry system is an open-circuit system that determines O_2_ consumption, CO_2_ production, and RER. Data were collected after 3 hr of adaptation in acclimated singly housed mice.

### Plasmids

Hairpins against *Dnmt1, Dnmt3a*, and *Dnmt3b* were purchased from Sigma. Lentiviral overexpression vectors for *Dnmt1, 3a*, and *3b* were subcloned into pCDH using various multicloning sites (XbaI/NotI for Dnmt1, EcoRI/NotI for Dnmt1, Dnmt3a, Dnmt3a-CM) and NotI sites, and hairpins targeting *Dnmt* transcripts were subcloned at *Age*I/*Eco*RI or purchased from Open Biosystems. Hairpin sequences are shown in **Supplementary Table 1**.

### Statistical Analysis

Data from independent experiments are presented as the mean ± SE. Statistical differences were assessed using PRISM4 software (GraphPad). Unpaired two-tailed Student’s t tests and two-way ANOVA were used. p < 0.05 was considered statistically significant.

**Supplemental Table 1.**
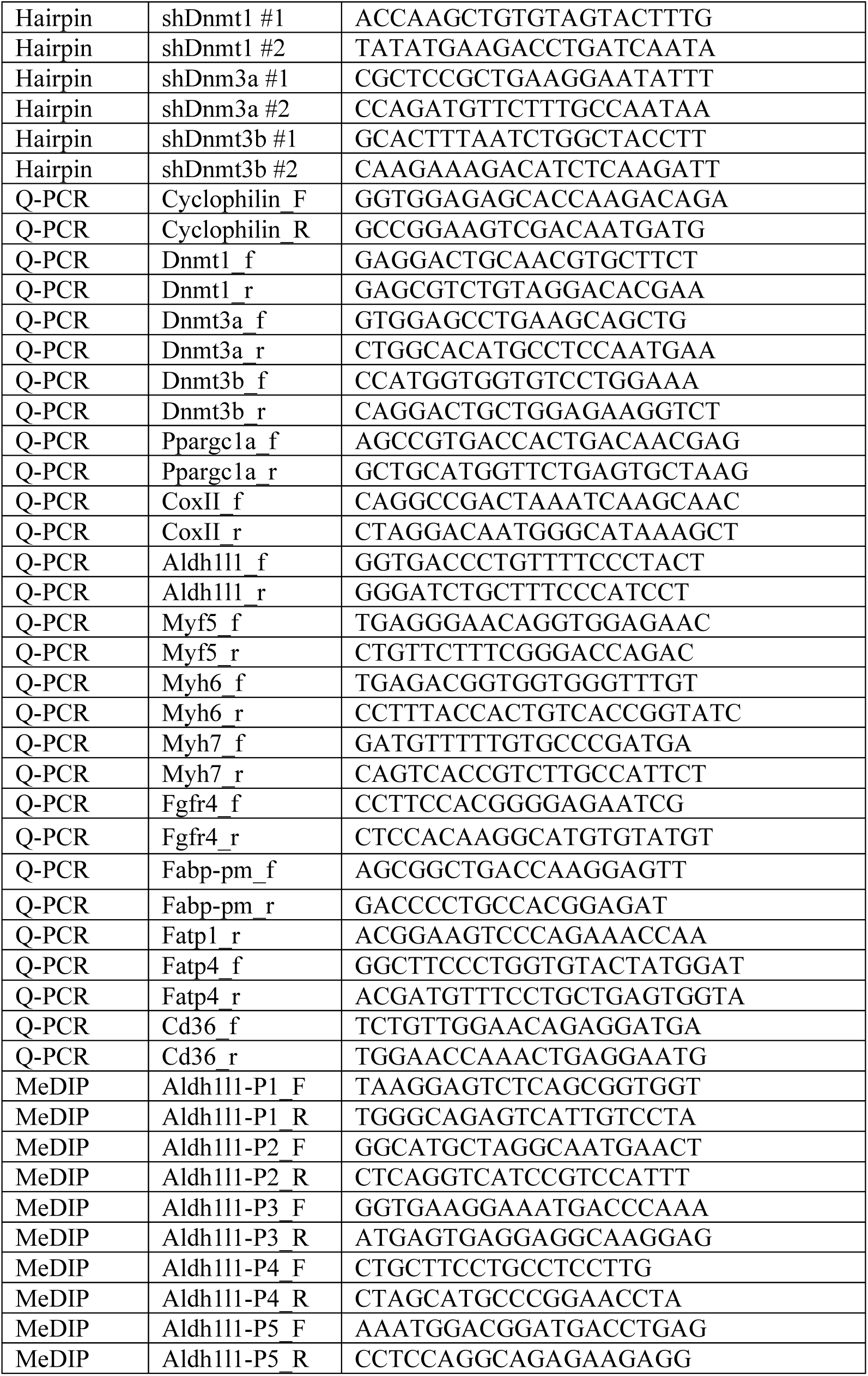
Oligonucleotide sequences used in this manuscript.

## Acknowledgements

We thank Dr. Hei Sook Sul, Dr. Jen-Chywan Wally Wang, and Dr. Anders Näär for helpful conversations about the manuscript. We are also grateful to Dr. Brian Black and Emily Wilson for technical help with histology.

## Author contributions

SK designed the experiments. Experiments were carried out by SDV, DY, JK, and SK. SK supervised all other experiments. SK and SDV wrote the manuscript.

## Funding

Work was funded by AHA Award # 19POST34380834 to DY and R01 DK116008 to SK.

## Competing interest

The authors declare no conflict of interest.

## Figure legends

**Supplemental Fig. 1.**
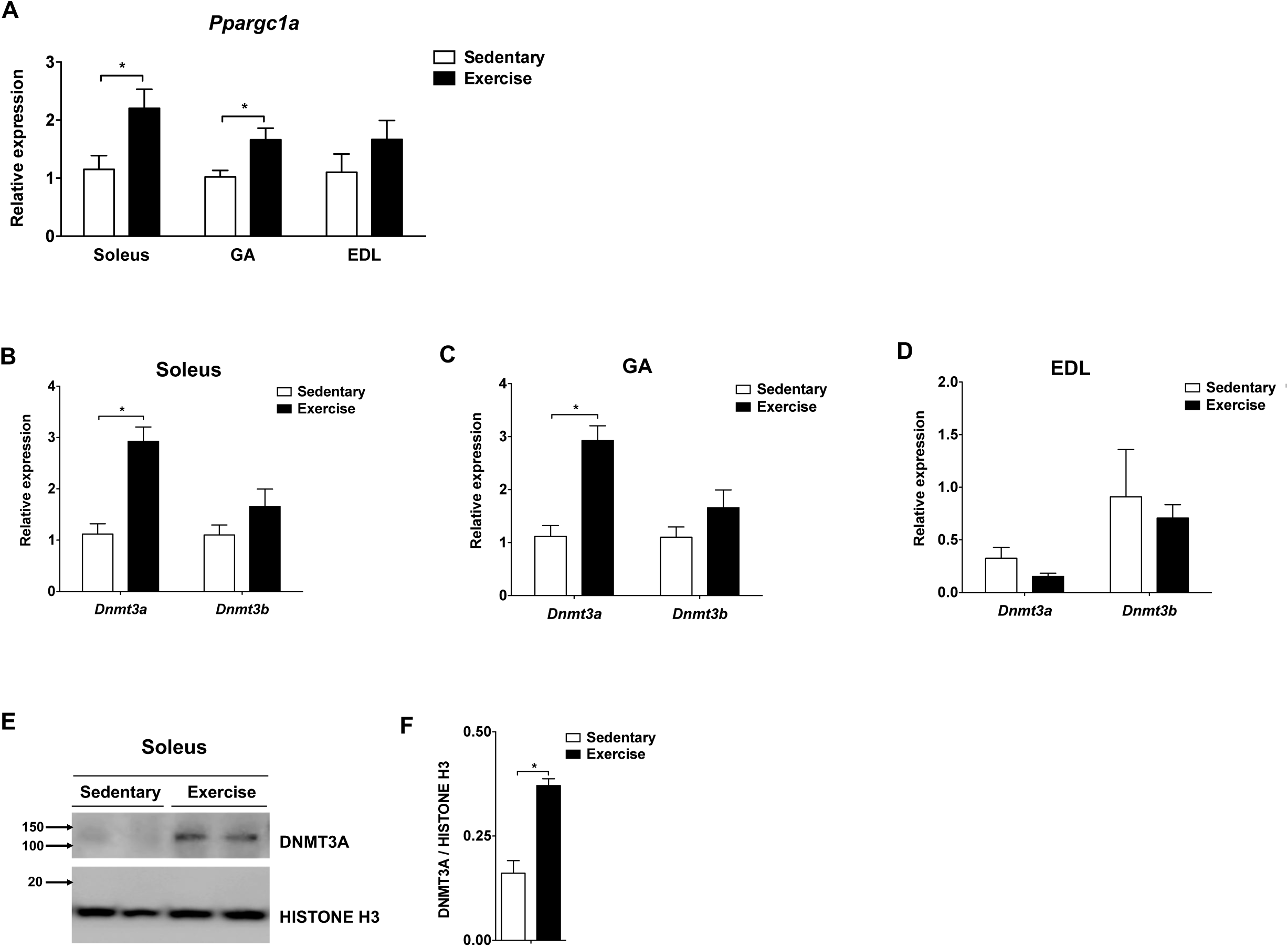
*Dnmt3a* levels are upregulated in soleus and GA muscles after a bout of endurance exercise. (**A**) *Ppargc1a* was measured in various muscle types from C57BL/6J mice at rest and after a bout of low-intensity exercise (*n* = 6, p < 0.05, Student’s *t*-test, mean ± s.e.m.), GA: Gastrocnemius; EDL: Extensor digitorum longus). (**B-D**) Transcript levels of genes encoding de novo DNMTs were measured in soleus (**B**), GA (**C**), and EDL (**D**) from C57BL/6J mice at rest and after a bout of low-intensity exercise (n = 6, p < 0.05, Student’s t-test, mean ± s.e.m.). (**E**) Soleus DNMT3A protein level in the nuclear fraction from **A** was measured by immunoblotting and (**F**) normalizing to Histone H3 using ImageJ (n = 2, p < 0.05, Student’s t-test, mean ± s.e.m.).

**Supplemental Fig. 2.**
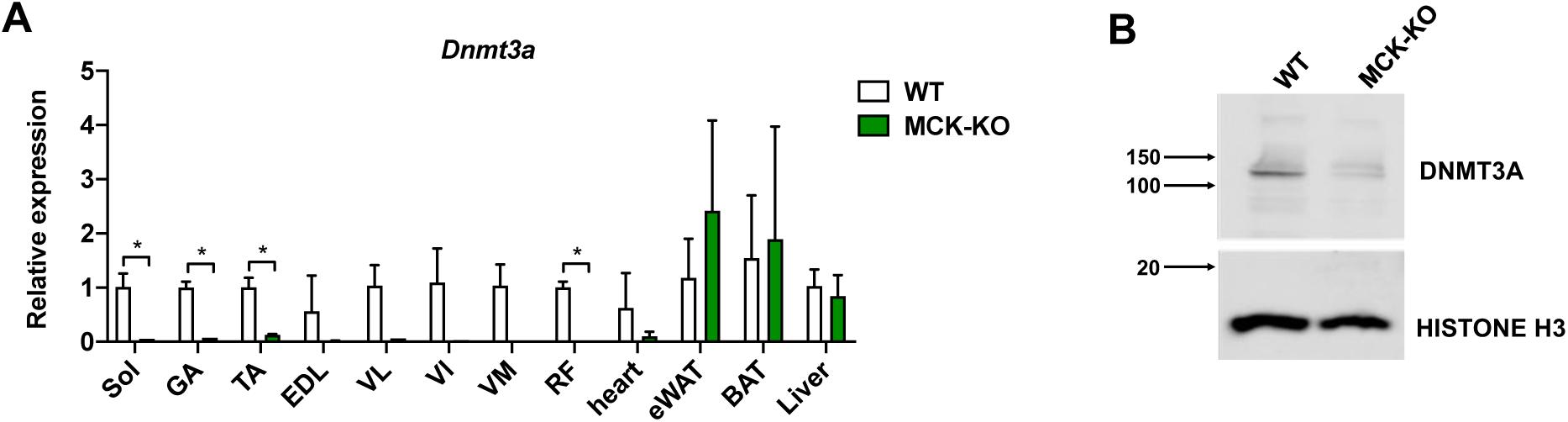
Establishment of muscle-specific knockout of *Dnmt3a* using MCK-Cre. (**A**) *Dnmt3a* mRNA expression was measured in various tissues from KO and WT mice by qPCR analysis (n = 5 mice, p < 0.05, Student’s *t*-test, mean ± s.e.m.). (**B**) DNMT3A protein expression was assessed by immunoblotting using nuclear extract from KO and WT soleus muscle.

**Supplemental Fig. 3.**
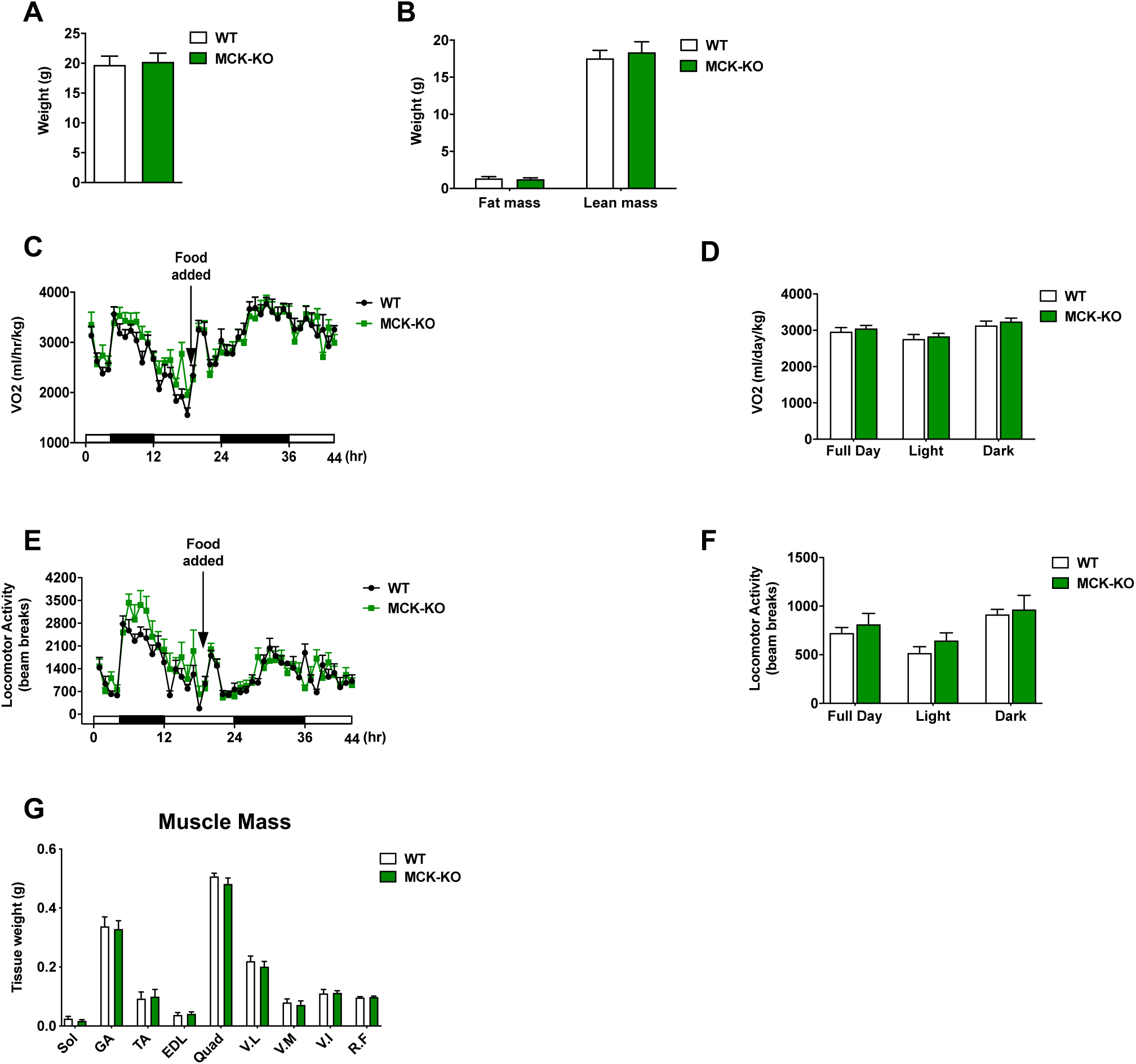
MCK-*Dnmt3a* KO mice do not show obvious change in energy homeostasis and muscle mass on chow diet. Body weight (**A**) and body composition (**B**) of MCK-*Dnmt3a* KO and WT mice on chow (n = 5 mice, p < 0.05, Student’s t-test, mean ± s.e.m.). Shown is the whole-body oxygen consumption rate (VO2, **C**) and averaged VO2 (**D**) of MCK-*Dnmt3a* KO and WT mice on chow during fasting (first 20 hrs) and fed (food added for next 25 hrs) conditions (*n* = 6 mice, p<0.05, p < 0.05, 2-way ANOVA, mean ± s.e.m.). (**E, F**) The whole-body locomotor activity of chow-fed MCK-*Dnmt3a* KO and WT mice during fasting (first 20 hrs) and fed (food added for next 25 hrs) conditions (*n* = 6 mice, p<0.05, p < 0.05, 2-way ANOVA, mean ± s.e.m. (**G**) Tissue weight of different skeletal muscle types from chow-fed WT and KO mice. (n = 5 mice, p < 0.05, Student’s t-test, mean ± s.e.m).

**Supplemental Fig. 4.**
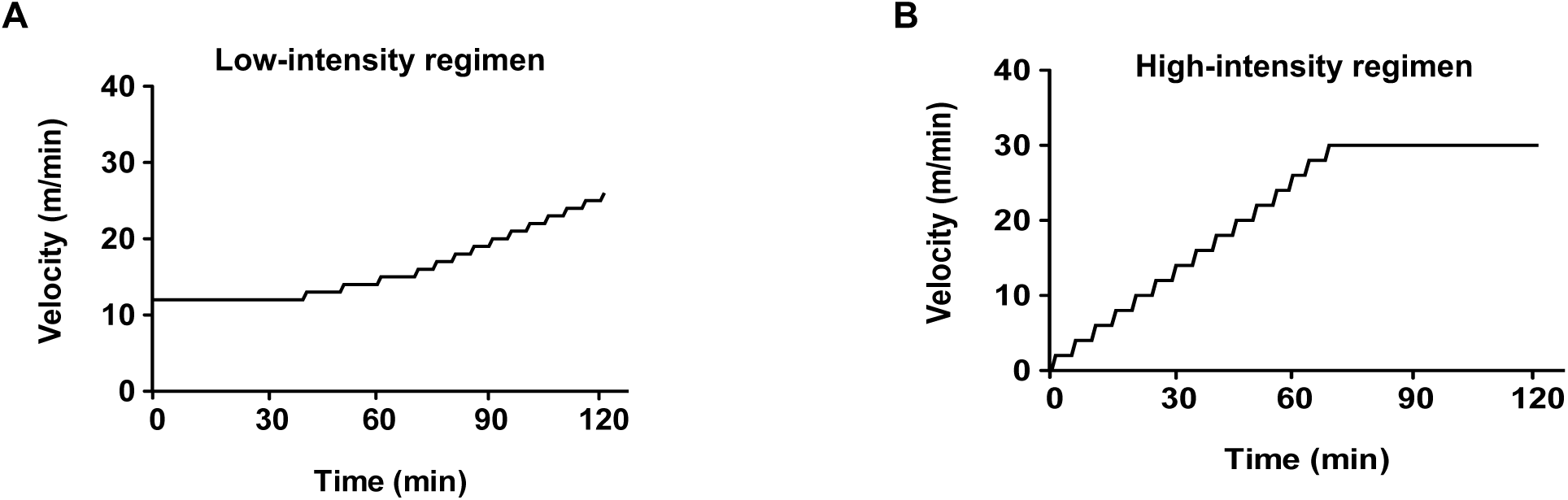
Schematic representation of exercise regimes. (**A**) Low intensity and (**B**) high intensity exercise regimens.

**Supplemental Fig. 5.**
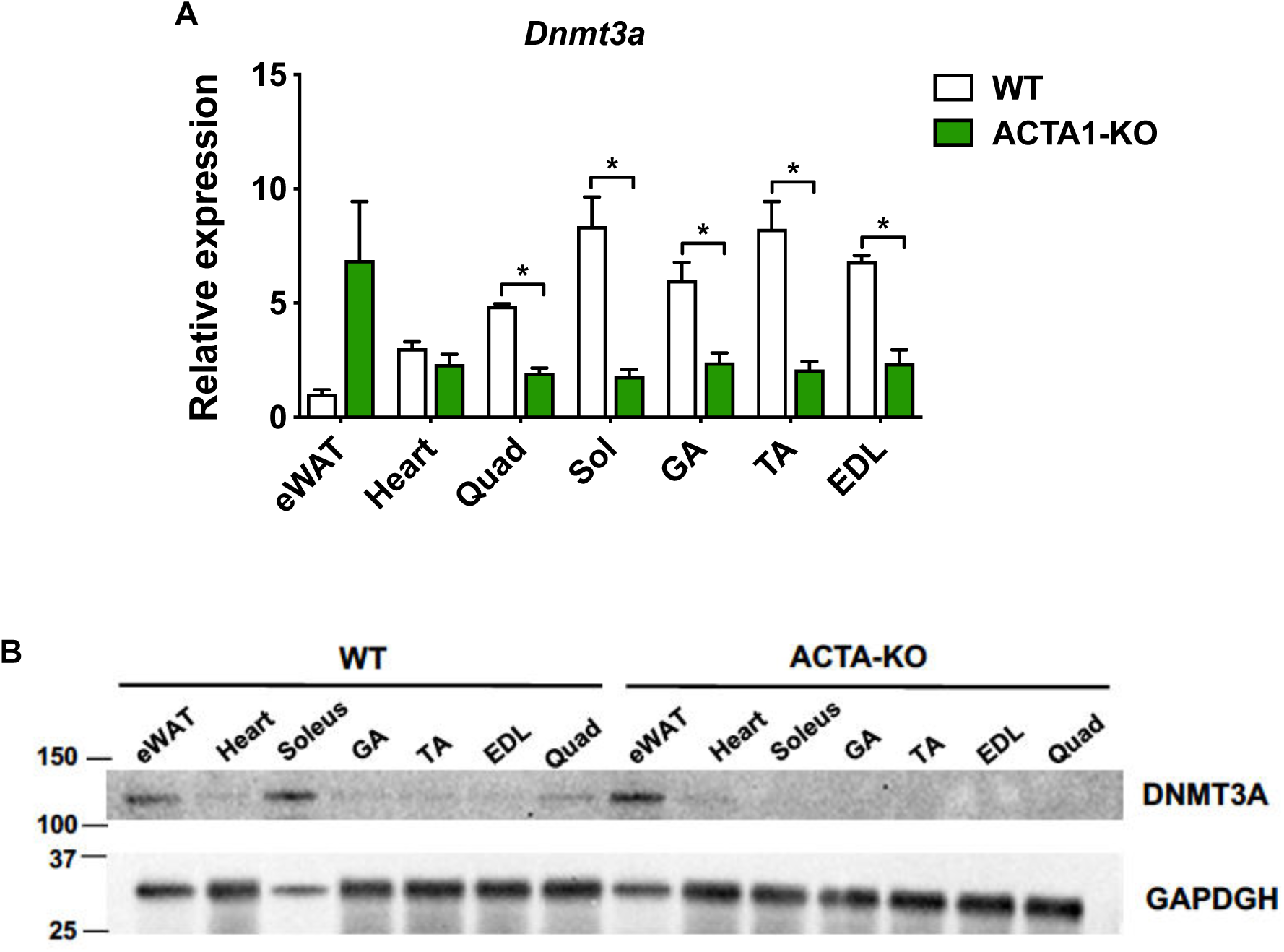
Establishment of muscle-specific knockout of *Dnmt3a* using ACTA1-Cre. (**A**) *Dnmt3a* mRNA expression was measured in various tissues from ACTA1-KO and WT mice by qPCR analysis (n = 5, p < 0.05, Student’s t-test, mean ± s.e.m.), (eWAT: epididymal white adipose tissue, Quad: Quadriceps, Sol: Soleus, GA: Gastrocnemius, TA: Tibialis anterior, EDL: Extensor digitorum longus). (**B**) DNMT3A protein expression was assessed by immunoblotting different tissue proteins from ACTA1-*Dnmt3a* KO and WT mice.

**Supplemental Fig. 6.**
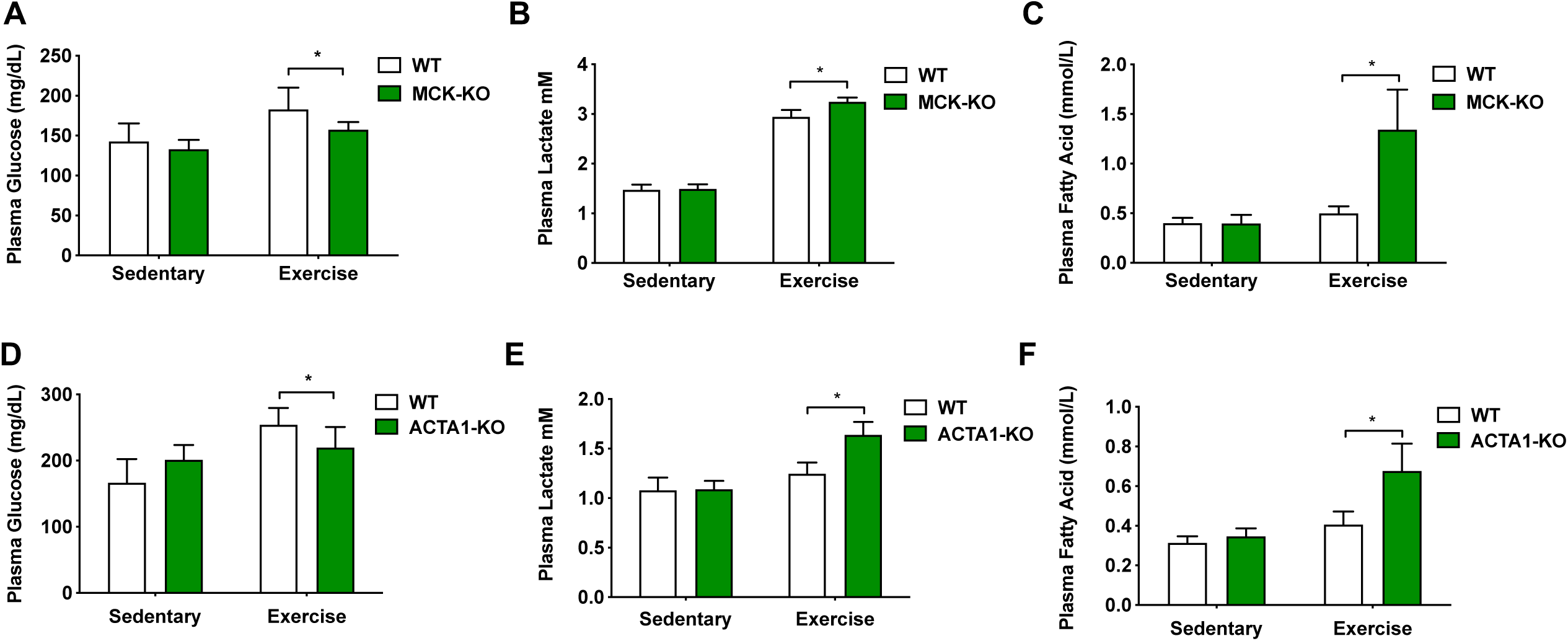
*Dnmt3a* KO mice display signs of an increased non-oxidative glycolysis upon exercise. (**A, D**) Plasma glucose levels were measured at sedentary and after a bout of low-intense exercise for 50 mins in MCK-*Dnmt3a* KO mice (**A**) and ACTA1-*Dnmt3a* KO mice (**D**) (*n* = 6, p<0.05, 2-way ANOVA, mean ± s.e.m.). (**B, E**) Serum levels of lactate were measured before and after a bout of low-intense exercise for 50 mins in MCK-*Dnmt3a* KO mice (**B**) and ACTA1-*Dnmt3a* KO mice (**E**) (*n* = 6, p<0.05, 2-way ANOVA, mean ± s.e.m.). (**C, F**) Serum levels of free fatty acid were measured before and after a bout of low-intense exercise for 50 mins in MCK-*Dnmt3a* KO mice (**C**) and ACTA1-*Dnmt3a* KO mice (**F**) (*n* = 6, p < 0.05, 2-way ANOVA, mean ± s.e.m.).

**Supplemental Fig. 7.**
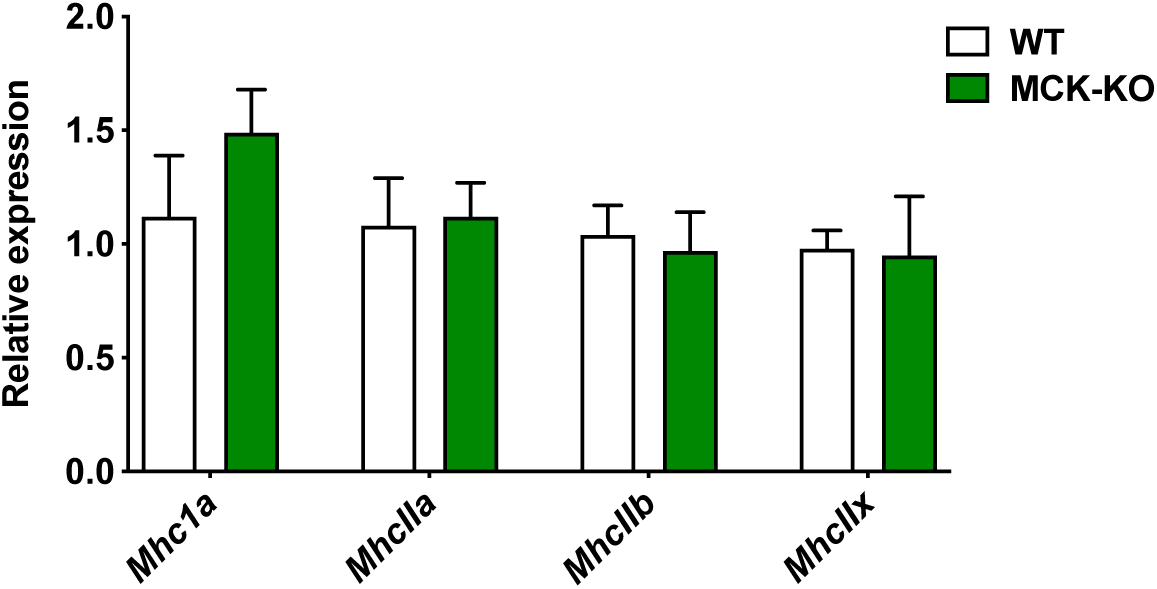
*Dnmt3a-*KO soleus muscle does not have transcriptional changes in fiber type– specific MHC isoforms. The mRNA expression of muscle subtype-specific MHC isoforms was measured in WT and KO soleus muscles (*n* = 5, p<0.05, Student’s *t*-test, mean ±s.e.m.).

**Supplemental Fig. 8.**
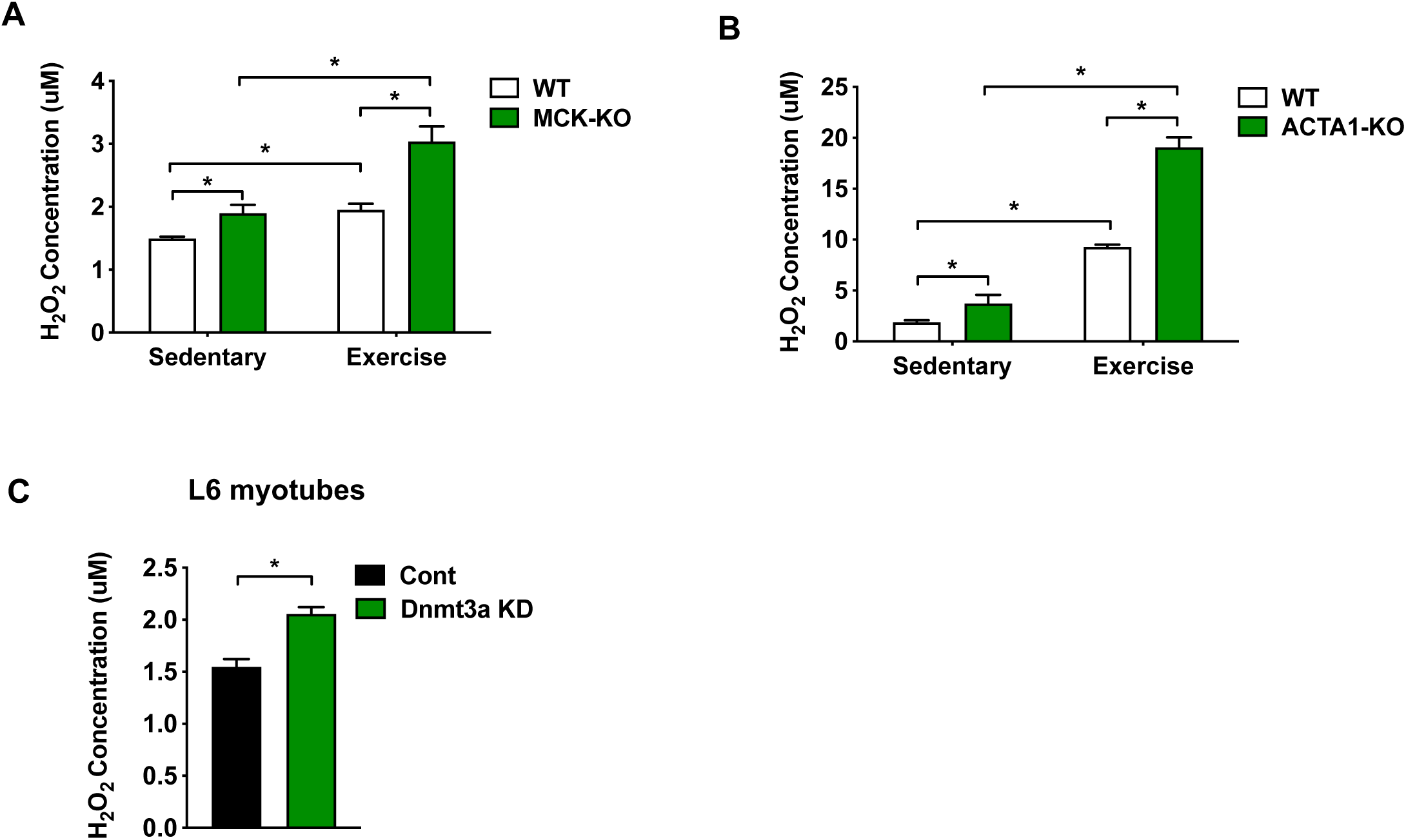
Increased ROS in *Dnmt3a-*KO soleus muscle and Dnmt3a knock-down myotubes. **(A-B)** Hydrogen peroxide (H_2_O_2_) levels were measured in WT and MCK-KO (**A**) and ACTA1-KO (**B**) at rest and after a bout of exercise (*n* = 6, p < 0.05, 2-way ANOVA, mean ± s.e.m.). (**C**) L6 myotubes were transduced with lentiviral expression plasmids for Flag-ALDH1L1 and GFP and H_2_O_2_ levels were measured from these cells (n = 6, p<0.05, Student’s *t*-test, mean ±s.e.m.).

**Supplemental Fig. 9.**
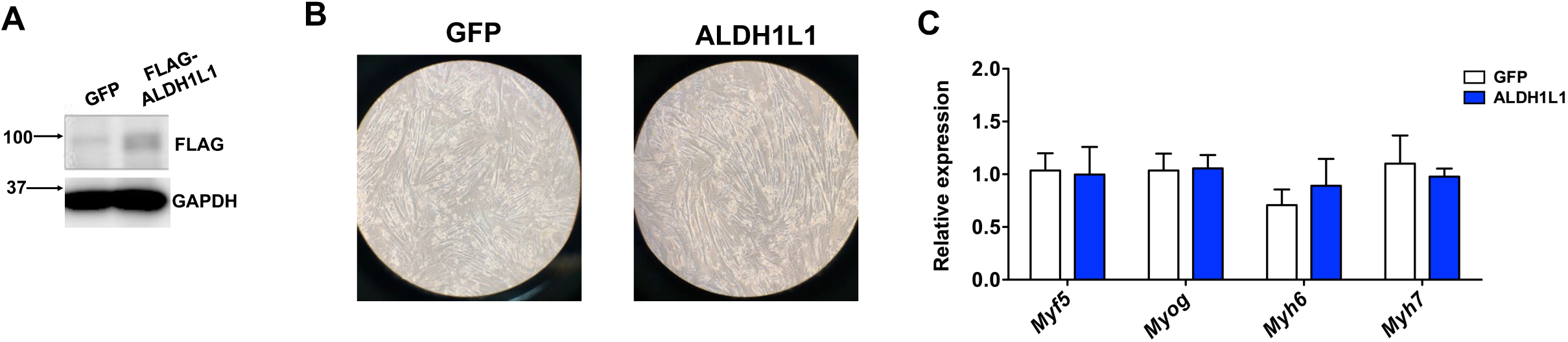
Overexpression of *Aldh1l1* does not affect differentiation status of myotubes. (**A**) L6 cells were transduced with lentiviral expression plasmids for Flag-ALDH1L1 and GFP. Flag western was performed to confirm the overexpression level of ALDH1L1. (**B**) Shown is the microscopic shot of L6 cells that overexpress Flag-ALDH1L1 and GFP from (**A**). (**C**) The mRNA expression of myogenesis markers were measured by qPCR in cells from (**A**). (*n* = 4, p<0.05, Student’s *t*-test, mean ±s.e.m.).

